# ShapeME: A tool and web front-end for de novo discovery of structural motifs underpinning protein-DNA interactions

**DOI:** 10.1101/2025.01.28.635290

**Authors:** Jeremy W. Schroeder, Michael B. Wolfe, Lydia Freddolino

**Author notes:** These authors contributed equally to this work (Jeremy W. Schroeder, Michael B. Wolfe).

## Abstract

Determining where transcriptional regulators bind within a genome is paramount to understanding how gene expression is regulated. Historically, position weight matrices (PWMs) have been used to define the binding preferences of DNA binding proteins^1^. However, PWMs treat the identity of each base in a sequence as an independent and additive measure of binding preference, which can limit their utility^2^. Models that consider higher order interactions between nearby bases yield greater success in predicting proteins’ binding to DNA, but for many proteins there is still substantial room for improvement in predicting and understanding the determinants of proteins’ binding to DNA^3^. In addition to DNA sequence motifs, structural motifs (e.g., a narrow minor groove width) are important determinants of binding for some DNA-binding proteins^4^. Despite the initial success of algorithms using structural features of DNA to predict binding properties of proteins from either ChIP-seq or SELEX data^5–8^, there remains a need for a *de novo* structural motif discovery framework which can be applied to data from a variety of experimental designs. Here, we present a unified workflow, capable of utilizing virtually any type of data representing sequence coverage or enrichment (e.g. ChIP-seq, RNA-seq, SELEX, etc.), to discover short structural motifs with explanatory power for a protein’s DNA binding preference. We couple the DNAshapeR algorithm^9^ with our own information-theoretic approach to *de novo* motif discovery, and wrap shape and sequence motif inference and model selection into a single tool called ShapeME. Application of our structural motif discovery algorithm to proteins with ChIP-seq data in ENCODE datasets reveals a subset of proteins where short structural motifs outperform the best PWM for that protein as determined from the JASPAR database, or as identified by the sequence motif elicitation tool STREME. Our approach offers a powerful and versatile framework for inferring structural DNA binding motifs, and will complement current sequence-based motif elicitation tools in discovery of protein-DNA interaction principles. A web-based interface to ShapeME is available at https://seq2fun.dcmb.med.umich.edu/shapeme, with full source code available at https://github.com/freddolino-lab/ShapeME.

## Introduction

Understanding and predicting interactions between DNA-binding proteins and the DNA sequences they target is fundamental to elucidating gene regulation in any organism. Traditionally, the Position Weight Matrix (PWM) has been used as a predictor and explanation for the sequence preferences of any particular DNA binding protein to great success^1^. However, PWMs treat the identity of each base in a particular sequence as an independent and additive measure for sequence preference, which can limit their utility for some transcription factor families^2^. More recent biophysically-motivated models have had greater success in predicting protein binding to DNA^10,11^, as have those that consider higher order interactions between each base^12,13^. Nevertheless, for many DNA binding proteins there is still substantial room for improvement in predicting and understanding the DNA target-based determinants of protein binding^3^. Recent studies have shown that some DNA binding proteins prefer more general structural elements, such as a narrow minor groove width, rather than a simple identification of the Watson-Crick face of the bases themselves^4^. Furthermore, the algorithm DNAshapeR has been developed to predict general DNA shape features from sequence information alone using all atom Monte-Carlo simulations^9^. Previous work has shown that using shape information together with one-hot encoded DNA sequence can predict the binding strength of several transcription factor families as determined by SELEX-seq^14^. An anchored approach, where DNA-shape features are calculated around a core PWM motif, has led to improvements in predicting the location of ChIP-seq peaks for many DNA binding proteins^5^ and the creation of a large database of shape profiles for motifs found in JASPAR and UniProbe^15,16^. However, the PWM-anchored approach will miss DNA shape patterns that are not easily captured by a PWM and its flanking region. Thus, de novo discovery of informative patterns in DNA shape parameters is necessary to fully realize the predictive contributions of DNA shape. Only a few approaches have been developed for the de novo discovery of short motifs in DNA shape information from either ChIP-seq^6,7,17^ or SELEX data^8^ and none are flexible enough to take advantage of many different data-types for which predictive motifs may be informative.

Here, we present a unified workflow, capable of utilizing either categorical data, e.g., ChIP-seq peaks vs. non-peaks, or continuous data, e.g., gene expression TPM values or ChIP-seq peak heights, to discover short DNA shape motifs with explanatory power for the binding preferences of DNA binding proteins. To accomplish this, we couple the DNAshapeR algorithm with our own information-theoretic approach inspired by the FIRE algorithm for discovering short sequence motifs^18^. Our algorithm scans the shapes in user-provided input data for seeds that have mutual information with the categories present in the input data (Fig. 1A). The list of informative seeds is pruned to a list of non-redundant seeds using conditional adjusted mutual information (CAMI). Remaining seeds are optimized using simulated annealing, followed by a second round of model pruning using CAMI in case distinct seeds converged to redundant motifs after optimization. Finally, LASSO regression is used to further retain the most informative remaining motifs.

**Figure 1.**
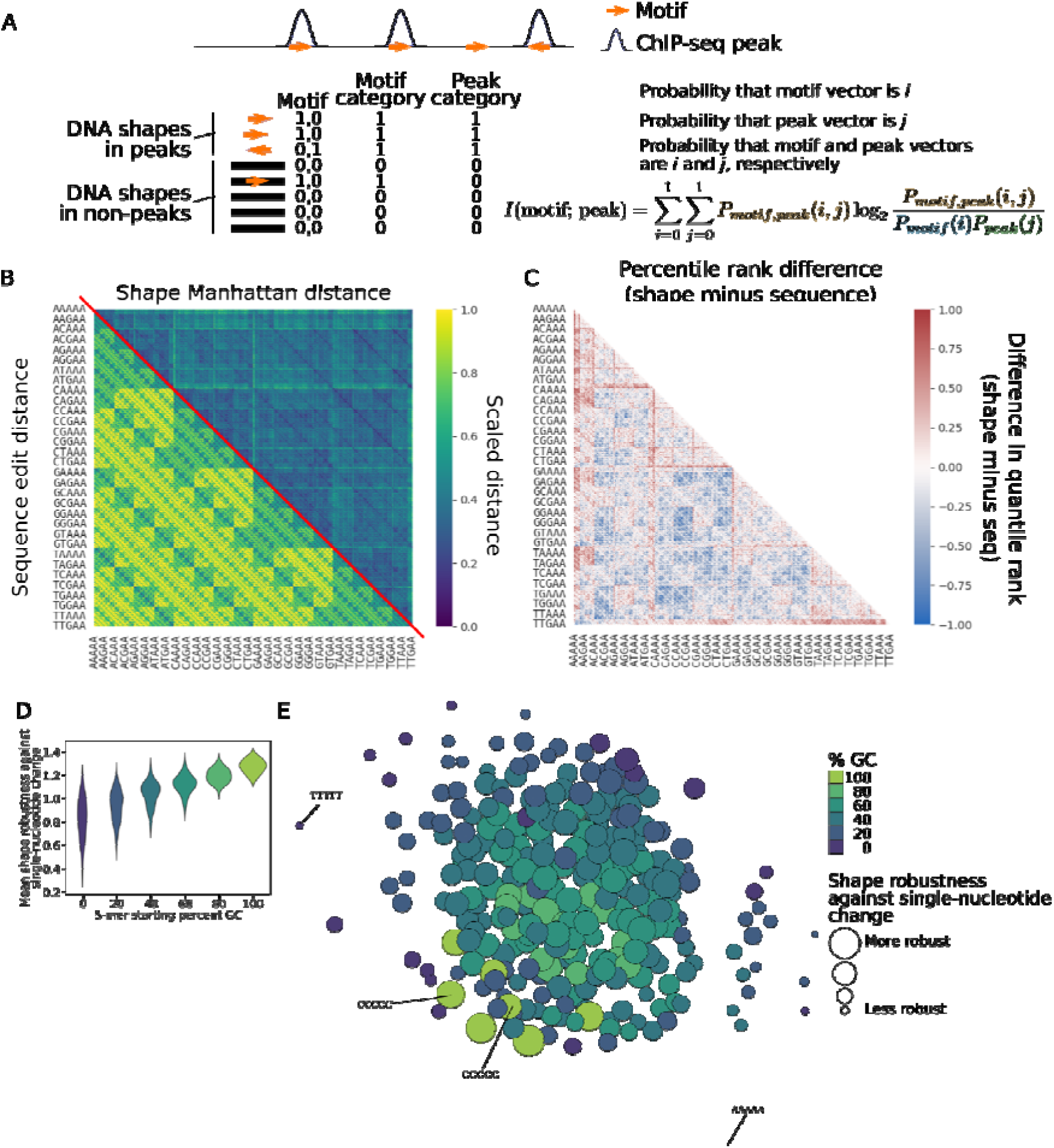
Using local DNA shapes and information theory for motif elicitation. A) A schematic depiction of ShapeME’s use of mutual information to quantify the extent to which motif presence informs input data scores. Note that for the sake of simplicity of presentation, the definition of mutual information is shown here, but ShapeME uses adjusted information (see methods). B) A heatmap displaying either the edit distance (lower triangle) or the manhattan distance (upper triangle) between all possible 5-mer sequences. Distances are scaled to a minimum of zero and a maximum of one in order to present them on a unified color scale. C) A heatmap demonstrating the difference in quantile rank between the shape distance rank and the sequence distance rank for each 5-mer. There are notable regions where shapes are consistently closer than sequence. D) 5-mer shape robustness at each GC content indicated in the x-axis. E) Graph representation of 5-mer shape robustness against single-nucleotide changes for 256 5-mers. Each node represents a 5-mer. GC content is encoded by node color and node size is proportional to the given 5-mer shape’s mean robustness (see methods for shape robustness definition) against all possible single-nucleotide substitutions. The weights of the edges between each node are the shape Manhattan distances between each 5-mer. AT-rich 5-mers are often near the exterior of the graph, whereas GC-rich 5-mers are closer to each other in shape space and have shapes which are more robust. The graph was plotted using Graphviz^45^ using the neato layout engine^46^.

The shape motif discovery method that we have developed is intended to complement currently available sequence motif inference tools through the identification of motifs in a new feature space, i.e., structural characteristics. As such, an important feature of our overall ShapeME workflow is that it also incorporates sequence motif inference using the popular tool STREME^19^. Sequence motifs identified by STREME are included into the overall motif model and are subject to CAMI-based filtering and LASSO regression along with the shape motifs present in the model. Therefore, ShapeME is able to automatically identify the most informative set of non-redundant motifs that inform protein-DNA interaction whether they are sequence or shape motifs.

To assess the performance of both DNA shape motifs identified by our algorithm, and the overall ShapeME workflow, we applied our method to a sample of 29 proteins with ChIP-seq data in the ENCODE portal and discover that shape motifs identified by the shape-based component of ShapeME often outperform both sequence motifs arrived at by STREME alone and the best PWM for that protein at the JASPAR database. In distinguishing true ChIP-seq binding sites from randomly sampled locations in the genome, of the 29 ENCODE datasets tested, 11 had shape motifs that outperformed sequence motifs, 12 had sequence motifs that outperformed shape motifs, and models considering either shape alone or sequence alone were both outperformed by models considering both shape and sequence for 15 cases. To take full advantage of the increased performance when both DNA shape and sequence are considered, ShapeME is able to natively perform motif inference using DNA shape and sequence, and will select the best set of non-redundant motifs to report to the user. We distribute ShapeME as an Apptainer container, but for most users we provide a simple web-based interface (https://seq2fun.dcmb.med.umich.edu/shapeme/) to infer motifs.

## Results

### DNA shapes provide a distinct landscape from DNA sequences

Building off of prior observations that DNA shape features can be highly informative for predicting the binding preferences of some proteins, we first sought to understand the extent to which the distance between short pieces of DNA in sequence space differed from the distance between DNA in local shape space. We thus generated all possible 5-mer DNA sequences, and placed a constant “AA” dinucleotide on each end of the 5-mers to enable shape prediction for the central 5 positions of each sequence of DNA. Qualitative assessment of all pairwise edit (for sequence space) or Manhattan (for shape space) distances revealed a dramatically different distance landscape for the two spaces (Fig. 1B). Edit and Manhattan distances were ranked and the difference in the quantile rank for each 5mer-to-5mer distance was calculated. Subtracting the quantile rank for a given edit distance from that of the comparison’s Manhattan distance showed that there are many regions of the distance landscape for which 5-mers are relatively more similar in shape space than their sequence space representations suggest, while the complement is true of other regions of the landscape (Fig. 1C). This finding implied that certain sequences may have underlying shapes that are more robust against changes in shape when a single-nucleotide substitution occurs.

We assessed the robustness (see Eq. 15 and associated methods) of each 5-mer shape against single-nucleotide changes. We found that shape robustness increased with increasing GC content (Fig. 1D-E). Our results suggest that to identify motifs that predict DNA-protein interaction sites, both DNA sequence and DNA shape must be considered in order to better capture the multidimensional similarities between DNA molecules that extend beyond a view of DNA as a simple sequence of nucleotides. In addition, we observe that the topology of the shape space is qualitatively different from that of sequence space, in that the nearest-neighbor sequences in one space are not necessarily close neighbors in another, which provides a major motivation for the consideration of shape-based motifs.

### ShapeME correctly identifies shape motifs in synthetic datasets

In order to test our ability to capture a “ground truth” shape motif in a variety of different contexts, we produced several synthetic datasets to assess the ability of ShapeME to infer motifs known to be present in input data. For the test datasets in this section, ShapeME was run in shape-only mode unless otherwise stated.

#### A single motif in every “positive” set sequence

We first tested the simplest test case, where a single target shape was placed into a subset of the sequences in a test dataset. We generated a set of 3000 random sequences, each 60 nucleotides long. We randomly assigned 20% of the sequences to the positive set. The randomly chosen sequence ACATGCAGTC was substituted at a random position on the plus strand of every sequence in the positive set, and into a randomly-selected 1% of sequences in the negative set. ShapeME was able to identify the shape motif, with an average 5-fold cross-validated area under the precision-recall curve (CV-AUPR) of 0.999 (Fig. 2A).

**Figure 2.**
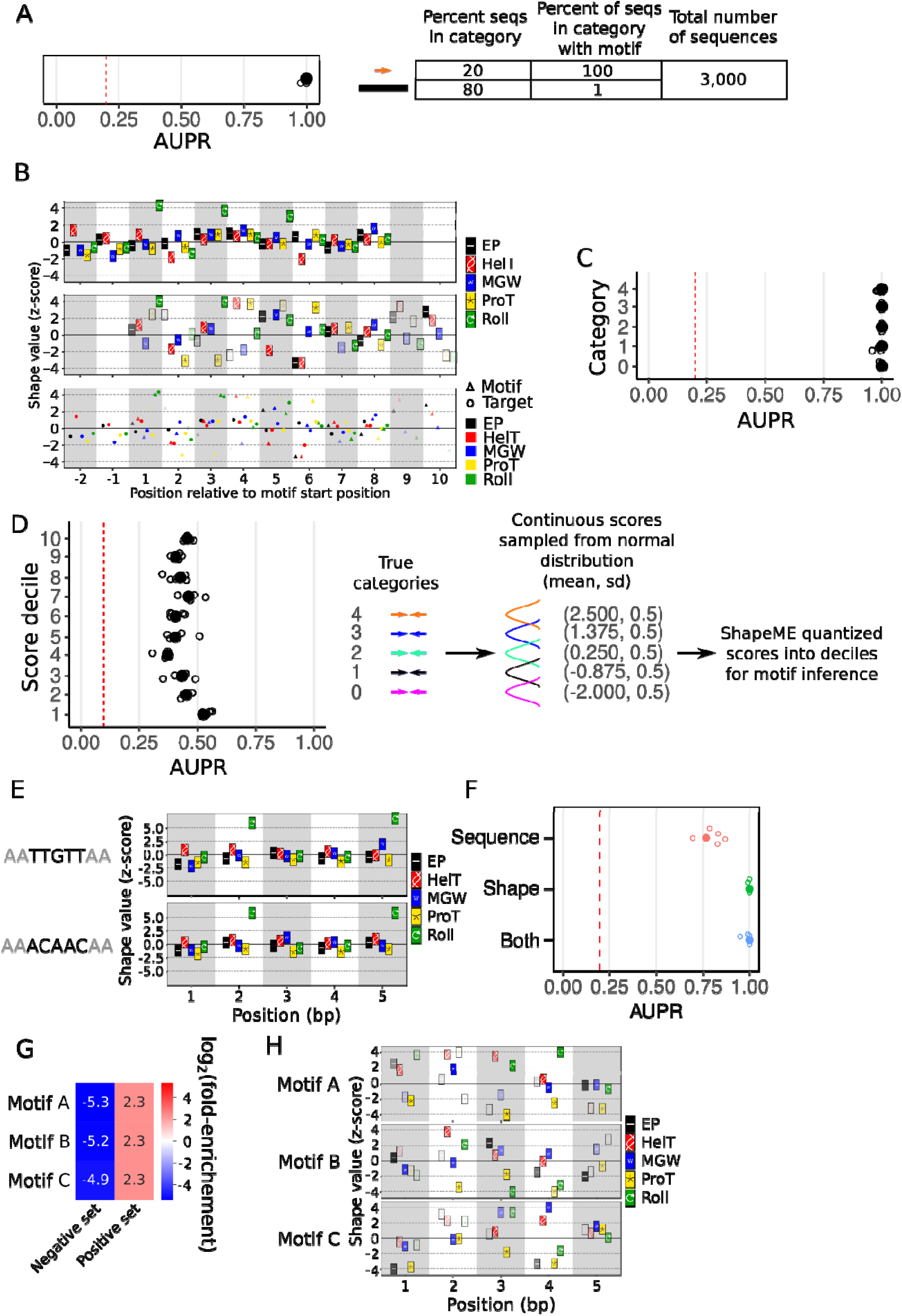
ShapeME identifies shape motifs in synthetic datasets. A) ShapeME performance when inferring a shape motif placed in binary input data in which a known target motif is present. For all such panels, filled circles indicate the AUPR for models trained and evaluated on the entire dataset, and smaller open circles indicate AUPRs for each fold of 5-fold cross validation. Note that several of the five open circles are often obscured by the larger filled circle. B) Top plot: target shape in the datasets in panel A. Middle plot: best-performing shape motif returned by ShapeME. Icon opacity denotes the weight applied to the given shape value at the position indicated by the x-axis. Bottom plot: aligned shape motif (triangles) and target shape (circles) plotted together. In the bottom plot, triangle opacity carries the same meaning as icon opacity for the shape motif logo in the middle plot. C) ShapeME performance on categorical input data with five categories. A separate known motif was placed once into each strand of all sequences in each category. D) ShapeME performance on a more difficult task. A separate target motif was placed into each strand of all sequences in each of five starting categories. Each sequence was assigned a continuous score via randomly sampling from a normal distribution, the mean of which depended on the motif present in the sequence (see schematic to the right). ShapeME was run using the continuous scores as inputs, and scores were quantized into deciles by ShapeME for motif inference. E) The shapes arising from each sequence selected to test the ability of ShapeME to identify shape motifs that converge from dissimilar sequences. The constant “AA” dinucleotides on each end of the selected sequences are shown in light gray, but the entire nine bases were inserted into the positive set of sequences to assure the desired shape 5-mer would result from sequence insertion. F-H) ShapeME results when run on input data containing the 9-mers indicated in paned E in the positive set of input sequences. F) Performance of ShapeME when run in each of its three modes. For all panels, vertical red dashed lines denote the AUPR expected by random chance. G) Heatmap demonstrating motif enrichments for each motif identified in sequence-and-shape mode. Only shape motifs were identified as informative. H) Motif logos for each motif identified by ShapeME in sequence-and-shape mode.

Recognizing the utility of PWMs for the visualization and interpretation of nucleic acid binding protein preferences, we sought to present ShapeME users with a shape logo that is as intuitively interpretable as commonly-used sequence motif logos. Our approach plots each of the five parameters ShapeME uses (electrostatic potential, EP; helical twist, HelT; minor groove width, MGW; propeller twist, ProT; roll, Roll) as icons with each icon positioned at its appropriate z-score value on the y-axis at each position on the x-axis, and with each position/shape combination’s weight encoded as the icon’s opacity, where a weight of zero would result in a transparent icon, denoting that the value of the shape parameter at the indicated position is unimportant (Fig. 2B).

Manual alignment of the target shape arising from the inserted sequence (Fig. 2B, top) with the shape motif inferred by ShapeME (Fig. 2B, middle) revealed the shape-based component of ShapeME is able to identify a known target shape motif in this synthetic dataset.

#### ShapeME identifies motifs in categorical input data

The case above represents binary input data, such as what one would generate when searching for motifs arising in ChIP-seq peaks, where peak DNA is in the positive set and randomly selected genomic DNA is in the negative set. However, it is often the case that a research question requires more refined binning of data than a simple binarization is able to provide. For example, one may desire to identify motifs enriched in promoters or enhancers driving expression of genes over a range of expression values. In this case, quantizing gene expression scores from, say, RNA polymerase ChIP-seq at promoters or perhaps transcript abundances from RNA-seq, into several bins may be useful. A major advantage of using mutual information to identify informative motifs is that it can be used naturally with categorical datasets, and is completely agnostic as to the origins of those categories, which could arise from genomic features, binning/clustering of one or more quantitative datasets, or any combination thereof.

To test the ability of ShapeME to recover known motifs in categorical data, we randomly generated 3000 sequences, each 60 nucleotides in length. We assigned each sequence to a category from 0-4, inclusive. We next generated five randomly selected known target sequences, and each target sequence was substituted into every one of its corresponding category of records in the test dataset as an inverted repeat. ShapeME identified shape motifs in the test dataset with high fidelity. The lowest AUPR for any fold of any of the five categories was 0.96 (Fig. 2C).

#### ShapeME quantizes continuous input scores to infer motifs

Rather than requiring users of ShapeME to manually categorize continuous scores, we wrote ShapeME to be able to quantize continuous input scores automatically. To test whether ShapeME could retrieve known motifs from continuous data which it had quantized into deciles (ShapeME allows users to choose how many bins to categorize continuous input data into), we generated two synthetic datasets: one with five categories of input sequences (Fig. 2D), and the other with ten categories of input sequences (Supplementary Fig. 1). Each category contained a distinct target motif on each strand. We then randomly sampled the scores associated with each input sequence from normal distributions, the centers of which depended on the motif in the given input sequence. Therefore, sequences containing equivalent target motifs were expected to be assigned similar input scores, but scores across sequences with differing target motifs will in many cases overlap (Fig. 2D, right}. Our goal was to simulate experiments in which the user is naive to how many true underlying categories are in the data, so while the input data had five ground truth categories for Fig. 2D, we had ShapeME quantize the continuous input scores into deciles, which is expected to yield motifs with lower performance than those learned in the two simple test datasets described above. As expected, the shape motifs inferred by ShapeME on this more difficult dataset achieved AUPRs near or slightly below 0.5 (Fig. 2D, left), which is significantly better than performance by random chance of 0.1.

For the dataset in Supplementary Fig. 1, the score sampling strategy used to generate the scores for each ground truth decile in the data were even more inter-mixed than in the dataset used for Fig. 2D (Supplementary Fig. 1A). It therefore presents a greater challenge for motif discovery, so we prepared datasets of varying size to demonstrate the effect of increasing dataset size on ShapeME’s power to detect true motifs present in input sequences. With only 2,000 sequences in the dataset, performance on this very challenging task was often little better than random chance (AUPR = 0.1), with an occasional fold during 5-fold cross validation achieving an AUPR greater than 0.4 (Supplementary Fig. 1B). As more sequences were added to this challenging dataset, performance of ShapeME both increased and became more consistent across folds (Supplementary Fig. 1B), with AUPRs near or exceeding 0.75 for many cases. Therefore, even on very challenging inference tasks, with sufficient data to inform motif discovery, ShapeME is able to identify informative shape motifs.

#### ShapeME identifies a single shape motif arising from two different sequences

Our final validation of ShapeME was performed by manually selecting two 5-mers from the set described in Fig. 1B-C. Our selection criteria were 1) the edit distance in sequence space must be 5, which is the maximum possible for two 5-mers, and 2) among 5-mers with edit distance equal to 5, the Manhattan distance in shape space must be minimized. The two sequences we selected were therefore ACAAC and TTGTT. We generated a synthetic dataset similar to that presented in the top plot of Fig. 2A, with the following modification: for the positive set of sequences we randomly chose either AAACAACAA or AATTGTTAA (note that the constant “AA” dinucleotide at each end is required to ensure the resulting central 5-mer shape is controlled) to place in the plus strand of the otherwise random DNA sequence (Fig. 2E). ShapeME can be run in one of three modes: sequence-only, shape-only, or both, i.e., both sequence and shape. Sequence-only mode uses the sequence motif elicitation tool STREME^19^, followed by motif pruning using CAMI (Eq. 4) and model selection using LASSO regression, the latter two steps being common between shape-only and sequence-only modes. “Both” mode infers informative shape motifs using our adjusted mutual information based approach, identifies candidate sequence motifs using STREME, performs pruning using CAMI and LASSO regression separately on each set of candidate motifs, merges the sequence and shape motifs into a single model, then performs a final round of LASSO regression for model selection. Therefore, “both” mode is able to select the best final set of motifs for the user, regardless of whether they are only sequence motifs, shape motifs, or a combination of the two types.

We ran ShapeME in each of its modes on the synthetic dataset arising from two distinct sequences that converge on a similar shape. With random performance on this dataset being approximately 0.19, each mode yielded informative motifs. On the full dataset, AUPRs for sequence, shape, and “both” modes were 0.77, 1.0, and 1.0, respectively. Sequence mode performance was more variable across folds during 5-fold cross validation than either shape or “both” modes (Fig. 2F). When ShapeME was run in “both” mode, no sequence motif was identified, whereas three shape motifs were identified, each enriched on a single strand in the positive set, as expected for this dataset (Fig. 2G).

Together, the better performance of shape-cognizant modes of ShapeME relative to its sequence-only mode, along with the more consistent performance across folds during 5-fold cross validation when ShapeME considered shape, demonstrate that when disparate sequences of DNA converge to a similar shape, as would often be expected for GC-rich DNA (Fig. 1D-E), not only is ShapeME able to identify shape motifs present in the data, but it is able to prune redundant, less informative motifs from the model when better-performing motifs providing similar information exist. Moreover, ShapeME is able to extract meaningful motifs from a variety of input data types.

### Shape motifs are often complementary to, and sometimes outperform, sequence motifs in biological datasets

Having established ShapeME as a useful tool to retrieve target motifs from synthetic datasets, we turned our attention to biological ChIP-seq data. We downloaded publicly available ChIP-seq data or ChIP-seq peak locations for several DNA-binding proteins to assess the determinants of DNA binding of each using ShapeME, with the working hypothesis that many DNA binding proteins (particularly those which show highly degenerate or low-performance sequence motifs) might primarily recognize DNA shape.

#### DNA binding by the bacterial nucleoid associated protein H-NS is associated with high roll

The bacterial nucleoid associated protein H-NS binds A/T-rich DNA, particularly when it contains TA steps, has a narrow MGW, and has low EP^20^. We downloaded H-NS ChIP-seq data for *E. coli* that were in either early-exponential (EE), mid-exponential (ME), transition (TS), or stationary (S) phase of growth. Because H-NS binds DNA broadly and forms extended filaments on DNA, we ran ShapeME on separate sets of input DNA sequences of either 100 bp or 300 bp in length. ShapeME also allows the user to choose the maximum number of shape motif occurrences on each strand of DNA, so we ran ShapeME on each input DNA length with a maximum hit count of either 1, 2, or 3. ShapeME identified informative shape motifs in all cases (Fig. 3A). Increasing input sequence length from 100 to 300 dramatically improved ShapeME performance for H-NS in all phases of growth, and increasing the maximum hit count on each strand provided improvements in performance at each input sequence length (Fig. 3A-B), consistent with the biological behavior of H-NS in forming extended oligomeric filaments on DNA. For the ShapeME run on 100 base-pair input DNA with a maximum hit count of 1, two shape motifs were retrieved. Each motif was specifically enriched unidirectionally, suggesting that H-NS either binds DNA directionally or forms filaments along DNA initially as a monomer (Fig. 3C). Furthermore, the logo for the more informative of the two motifs is punctuated by positions with elevated roll (Fig. 3D, Motif A, positions 1, 3, 5, 8, and 10). We therefore suggest that the statistical association between TA steps, which cause increased roll^21^, and H-NS binding identified in prior work^20^, may have indirectly identified the association between high roll values and H-NS binding.

**Figure 3.**
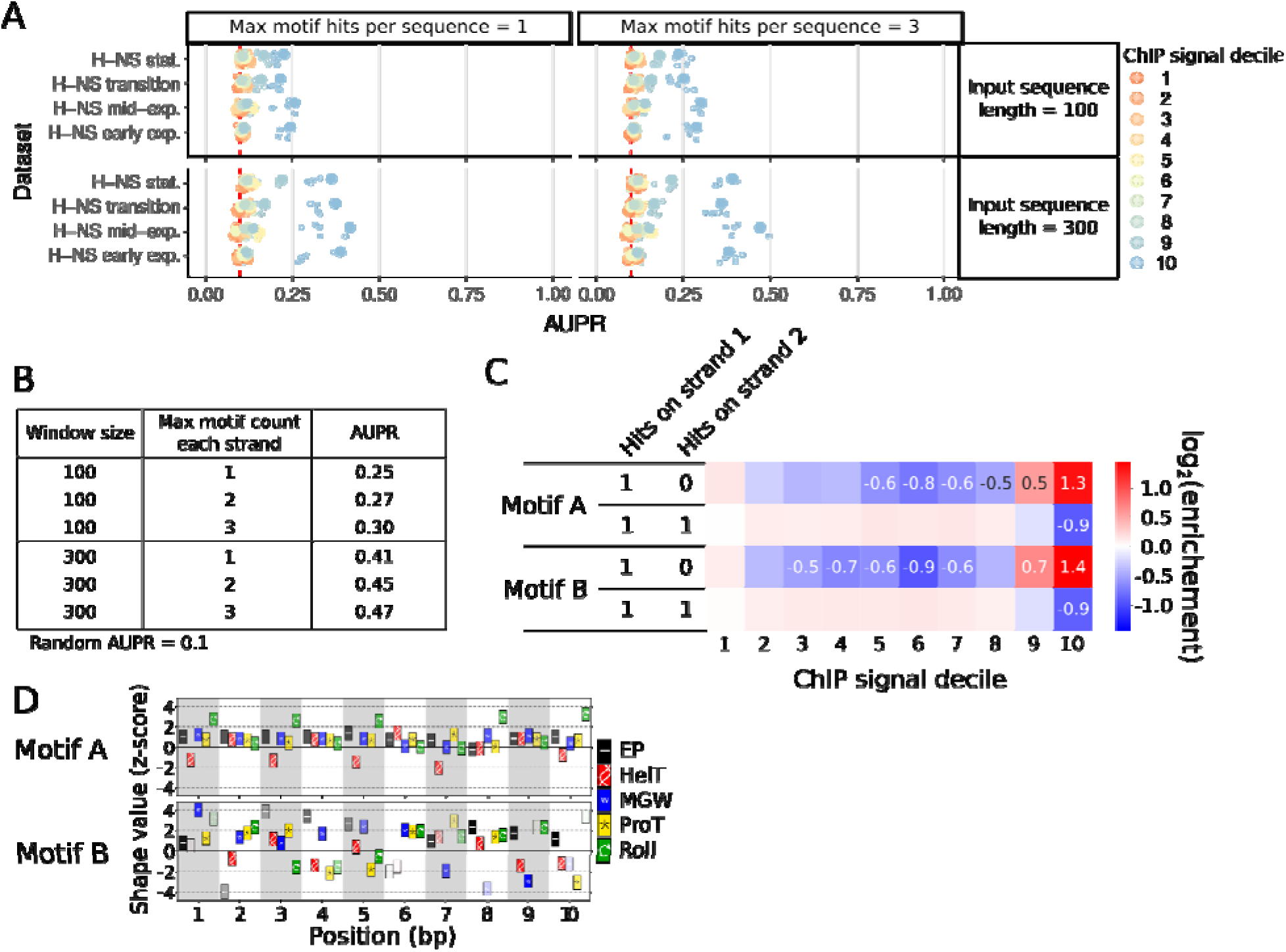
ShapeME identifies shape motifs in ChIP data for the bacterial nucleoid associated protein H-NS. A) Performance of ShapeME in shape-only mode when run on H-NS ChIP-seq data with the indicated input sequence lengths and maximum allowed motif occurrences. Filled circles are the performance on the full dataset and smaller open circles represent performance on each of 5 folds prepared for 5-fold cross validation. Random performance is indicated by the vertical red dashed line in each subplot. B) ShapeME performance in all runs using the N-NS mid-exp data. A baseline predictor would give an AUPR of 0.1 in all cases. C) Enrichment of each of the two shape motifs identified when ShapeME was run on 300 bp input sequences with the maximum motif count on each strand set to 1. D) Logos for each motif identified using 300 bp input sequences with the maximum motif count on each strand set to 1.

#### Shape motifs complement sequence motifs to predict interaction sites for many human transcription factors

We tested the ability of ShapeME to identify motifs in binding sites for a compendium of human transcription factors. We downloaded the locations of peaks passing the irreproducible discovery rate threshold from the ENCODE project^22–25^ for each of 29 human transcription factors (see Table 3 for details). For datasets with greater than 1,000 peaks, we randomly selected 1,000 peaks to include in the analysis. By applying ShapeME in shape-only mode, sequence-only mode, or “both” mode we were able to establish which of the selected proteins relied primarily on local shape, sequence, or a combination of the two to inform their binding to DNA. Additionally, we compared ShapeME results to those achieved using the sequence motif at the JASPAR database as a known sequence motif^26^. ShapeME evaluated the performance of the JASPAR motif alone, or was run in “both” mode, bypassing the STREME step in favor of using the JASPAR motif as the sequence motif. ShapeME then performed model selection as usual when run in “both” mode. Therefore, ShapeME was able to either integrate the JASPAR motif and additional shape motifs it identified into a single model, or to choose one type of motif to return as the best model for predicting protein-DNA interaction sites. A subset of the 29 ChIP-seq datasets we ran is presented in Fig. 4, and the full set is shown in Supplementary Figure 2.

**Figure 4.**
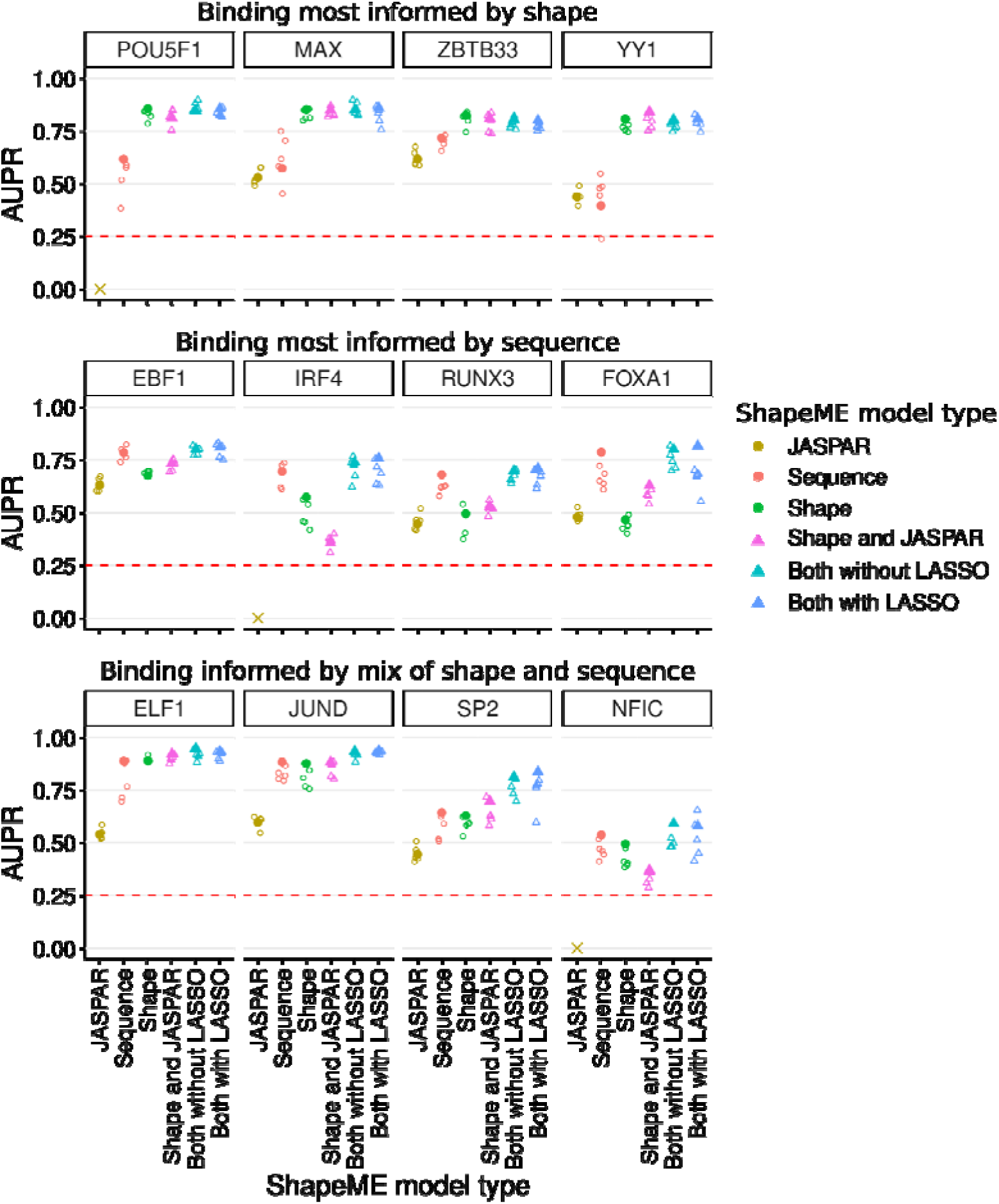
Shape motifs consistently outperform sequence motifs on ENCODE datasets. AUPRs are presented for the ENCODE data for the protein indicated above each plot. The “Both without LASSO” points represent motif models arrived at by running ShapeME in “both” mode, but leaving out the model selection step of LASSO regression. For detailed ShapeME performance on all ENCODE datasets used in this work, see Supplementary Figure 3. The “X” symbol for POU5F1, IRF4, and NFIC indicates that the motif at the JASPAR database, MA1115.1, MA1419.1, and MA0161.2, respectively, were not informative on the ENCODE data used here.

We were able to bin transcription factors into three categories based on the relative performance of shape and sequence motifs in predicting their binding profiles: 1) binding was most informed by DNA shape, as indicated by shape mode having the best performance (Fig. 4, top), 2) binding was most informed by sequence (Fig. 4, middle), or 3) both sequence and shape were required by ShapeME to achieve best performance (Fig. 4, bottom). While we show a balanced sampling across these three categories in Figure 4, in practice, purely shape-based motifs significantly outperformed JASPAR motifs in 22, and achieved rough parity in 2, out of 29 cases. Shape-based motifs significantly outperformed newly inferred sequence-based motifs in 12, and achieved rough parity in 5, of 29 cases. And, in an example of the utility of ShapeME’s ability to report a combination of sequence- and shape-based motifs, hybrid motifs incorporating both shape and sequence information provided significant improvement over newly inferred sequence-based motifs in 23, and achieved rough parity in the remaining 6, of 29 cases (see methods for details). The most extreme example of shape-dominated binding was YY1 – for this TF, shape-only mode performed far better (AUPR; 0.81) than either the JASPAR motif or sequence-only mode (AUPRs; 0.44 and 0.40, respectively). Inclusion of either the JASPAR motif or motifs identified by STREME did not appreciably improve performance beyond shape motifs alone (AUPRs; 0.84 and 0.81, respectively). The near opposite can be stated of ShapeME results on FOXA1 binding sites, for which the sequence motifs identified in sequence-only mode achieved an AUPR of 0.79, and addition of shape information did not appreciably improve performance (Fig. 4, middle-right sub-plot).

We also identified transcription factors for which the sequence and shape information considered together in “both” mode surpassed performance of either sequence information or shape information alone. For example, DNA binding sites for JUND, a member of the AP-1 transcription factor family, were well-predicted by ShapeME results for sequence-only and shape-only modes, each with an AUPR of 0.88. However, “both” mode achieved an AUPR of 0.94 (Fig. 4, bottom).

Together, these ShapeME results on biological datasets reveal that both sequence and shape information should be considered when attempting to characterize determinants of protein-DNA interaction. A major benefit of using ShapeME is that it can automatically select, among several candidate sequence and shape motifs, the most informative set of non-redundant motifs to report to the user.

### ShapeME outperforms the alternative shape motif inference tool ShapeMotifEM

As discussed above, other shape motif inference tools exist, many with the caveat that shape motif discovery is seeded by previously identified sequence motifs^15,16^ or that they are useful for identifying motifs comprising only a single shape parameter and cannot generalize to motifs which are composite of several shape parameters^6^. Recently ShapeMotifEM was published, which performs *de novo* shape motif discovery and generalizes to multiple shape parameters^17^. We compared performance of ShapeME to ShapeMotifEM on the human transcription factor ENCODE data used above, but filtering to retain the peaks with signal values exceeding the 95th percentile of all binding scores. Running ShapeME (in shape-only mode) and ShapeMotifEM on these datasets revealed that ShapeME motifs outperform motifs returned by ShapeMotifEM in all cases that ShapeMotifEM was able to run to completion (Fig. 5), and in 59% of cases ShapeMotifEM crashed without reporting results.

**Figure 5.**
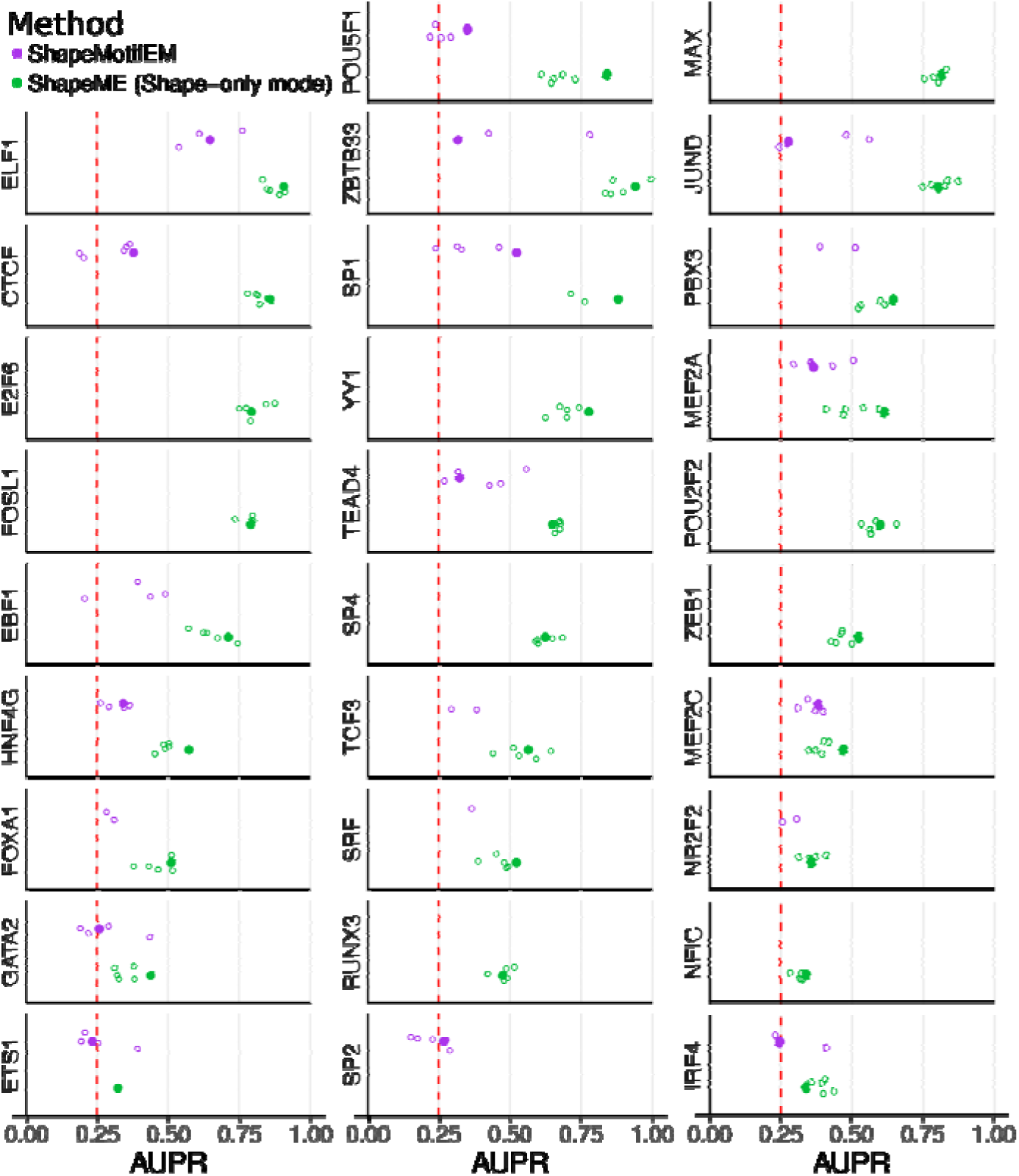
ShapeME outperforms ShapeMotifEM in benchmarks using ENCODE data. ShapeME and ShapeMotifEM performance for all peaks exceeding the 95th percentile signal strength in ENCODE data. For each dataset, three times as many random genomic sequences were present in the dataset than peak sequences. AUPRs are presented for ShapeME run in shape-only mode and ShapeMotifEM for each protein indicated to the left of each subplot. Large points indicate the AUPR resulting from training and evaluating motif models using all data, and smaller open circles denote AUPRs from each of five folds of the data used for 5-fold cross validation. Missing points or circles in ShapeME results are due to no informative shape motif having been identified. Missing points or circles in ShapeMotifEM results are due to either no motif being identified or an error during the ShapeMotifEM run. The red dashed line indicates the expected background performance in each case for a random predictor.

### ShapeME run on RNA-seq results reveals potential regulatory hierarchy of JunD

Inspired by the performance of ShapeME on JunD ChIP-seq data (Fig. 4, bottom), and because ShapeME was designed to be generally useful on a variety of input data types, we next tested whether ShapeME would retrieve JunD motifs (Fig. 6A) from enhancers associated with differential expression of nearby genes as observed during CRISPRi-mediated knock down of JUND. We performed a differential expression analysis of JUND knock down vs. a control guide RNA using publicly available data (see Methods for details). The input sequences were taken to be the central 100 bp of the closest enhancer to each gene. Any gene with no enhancer within 450 kbp was dropped from the analysis. For genes with multiple enhancers within the gene, the first enhancer (in chromosome coordinates) was selected for the analysis. We first ran ShapeME on the described sequences where each sequence was assigned a value of zero if the shrunken log_2_(fold-change) due to JUND knock down was negative, or one if it was greater than or equal to zero. ShapeME was run in “both” mode and returned a single shape motif, which was enriched unidirectionally in enhancers near up-regulated genes and bidirectionally in enhancers near down-regulated genes (Fig. 6B).

**Figure 6.**
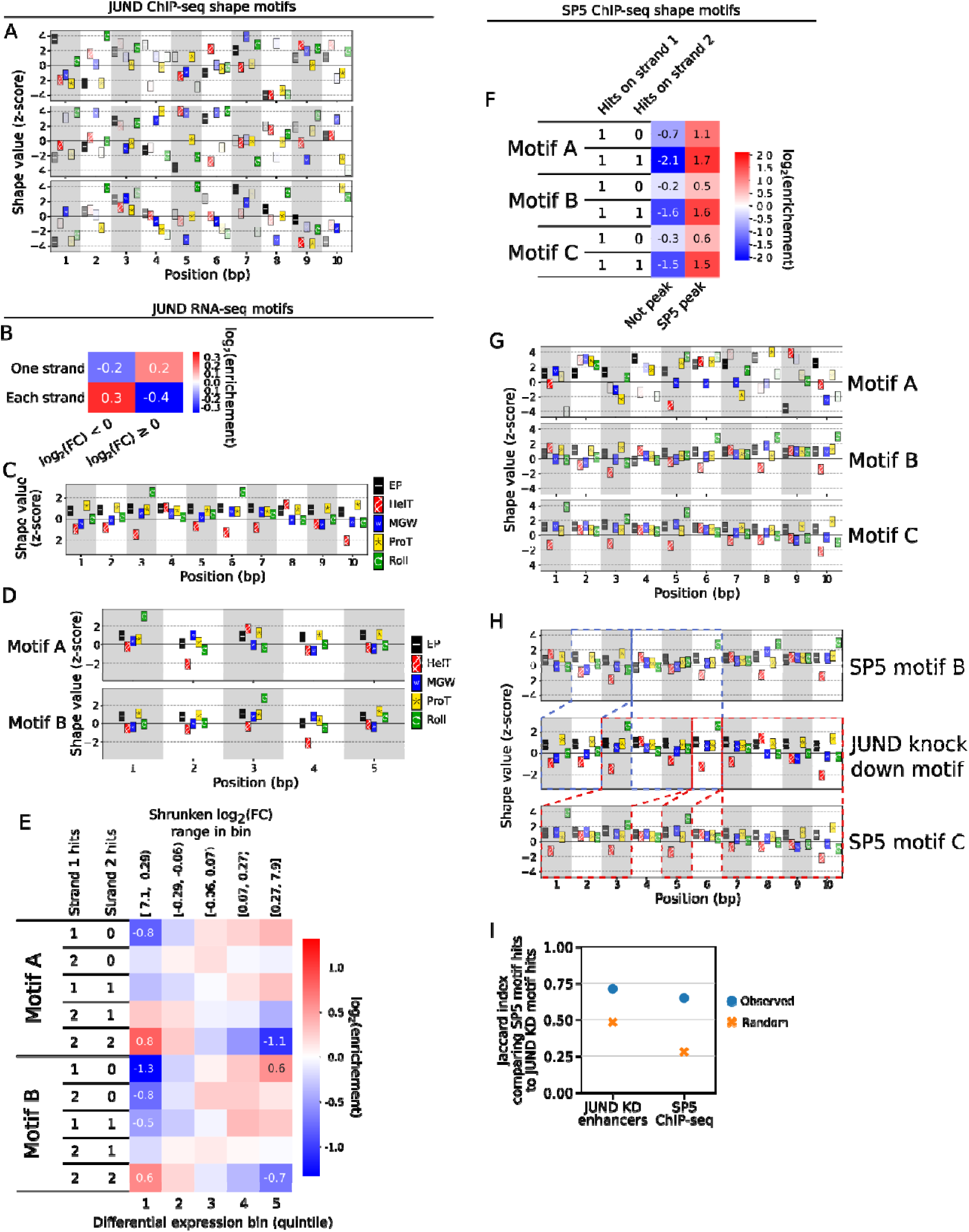
Identification of SP5 as potential factor explaining changes in gene expression upon JUND knockdown. A) Logos representing the three shape motifs returned by ShapeME run in “both” mode on JunD ChIP-seq data. B-E) ShapeME results using RNA-seq data from JUND knock down. The closest enhancer to each gene was used as input sequences and shrunken log_2_(JUND KD/control guide RNA) as input scores for each enhancer. B) Result of running ShapeME in “both” mode. A single shape motif was enriched in the enhancers nearest to genes with increased expression when it matched a single strand, and enriched in enhancers nearest to genes with decreased expression when it matched on both strands. C) The 10-mer shape motif logo enriched in enhancers associated with JUND knock down. D) Shape motif logos for both motifs yielded by ShapeME when allowing ShapeME to quantize continuous shrunken log_2_(JUND KD/control guide RNA) input scores into quintiles, setting the maximum number of hits on each sequence to 2, and searching for 5-mer shape motifs. E) Enrichment of each shape motif in the lowest quintile enhancers specifically when they hit each strand twice, and in the top quintile enhancers when they hit either a single or both strands once. F) Enrichments of 10-mer shape motifs in SP5 peaks. G) Shape motif logos for SP5 binding. H) Shape motif logos for SP5 motifs B and C manually aligned to the JUND KD motif presented in panel C. Local regions of high similarity between the JUND KD motif and SP5 motifs B and C are represented by dashed blue and red outlines, respectively. I) Jaccard indices for overlap between motifs trained on JUND KD enhancers and SP5 ChIP-seq data. Indices were calculated for each dataset indicated in the x-axis. The “random” value is the mean of 100 random selections of hits from the indicated dataset for each set of motifs.

Visual inspection of the shape motif logo suggested the 10-mer may have represented a repeated underlying 5-mer motif (Fig. 6C), so we re-ran ShapeME, this time directing it to search for 5-mer shape motifs and setting the maximum allowable motif hits per strand to two. In addition, for this second ShapeME run we allowed ShapeME to quantize the continuous shrunken log_2_(fold-change) values associated with each enhancer into quintiles. Indeed, this second ShapeME run yielded two qualitatively similar motifs (Fig. 6D) both of which were most strongly enriched in enhancers near the most downregulated genes, specifically when they appeared on both strands twice (Fig. 6E). However, when either motif hit a single strand or both strands only once, they were enriched in enhancers near upregulated genes (Fig. 6E). We discuss potential implications of this observation at the end of this section. While the 5-mer motifs (Fig. 6D) both matched well to segments of the 10-mer motif previously identified (Fig. 6C), none of the motifs identified using our RNA-seq analysis matched well to the JunD ChIP-seq motif (Fig. 6A). We therefore sought to determine whether the motifs identified using RNA-seq data from JUND knock down might be pointing toward the activity of a separate factor, potentially regulated by JunD.

For sequence motifs, after a motif of interest has been identified a next step is often to use TomTom to infer a set of proteins that may be responsible for the motif’s existence in a dataset^27^. While there is not yet a TomTom analog for shape motifs, we developed a companion tool to ShapeME called <u>Shape</u>-motif Instance <u>T</u>ool (ShapeIT) to identify instances of a shape motif in query DNA. ShapeIT may be thought of as the shape analog of FIMO from the MEME suite ecosystem^28^. As an interim analog to TomTom, we searched for TFs that would match the binding sites identified for the potential JunD-regulated factor: we ran ShapeIT to identify each instance of the 10-mer shape motif presented in Fig. 6B in the enhancer sequences used for motif elicitation and extracted the 10-mer nucleotide sequence underlying each shape motif instance. MEME was then used to identify sequence motifs in the resulting sequences^29^, and the sequence motifs returned by MEME were used as queries to TomTom. Among the results returned by TomTom was the transcription factor SP5, which is slightly down-regulated due to JUND knock down, with a shrunken log_2_(FC) = -0.56 and a q-value = 0.034. We reasoned that the shape motif revealed by ShapeME run on JUND knock down RNA-seq data could be an SP5 motif.

To test whether SP5 could be the factor driving shape motif identification in JUND knock down RNA-seq data, we downloaded SP5 ChIP-seq peaks from ENCODE and ran ShapeME to search for 10-mer shape motifs enriched in SP5 binding sites. Three shape motifs were identified as enriched in SP5 peaks (Fig. 6F-G), two of which (Motifs B and C in Fig. 6H) were qualitatively very similar to the 10-mer shape motif retrieved from enhancers near down-regulated genes during JUND knock down (Fig. 6B-C), with several positions having elevated Roll and somewhat low HelT (Fig. 6F-H). Motifs trained on JUND knock down RNA-seq results and those trained on SP5 ChIP-seq peaks identified similar sequences as hits when ShapeIT was run to identify motif instances in either the enhancer sequences or the SP5 peak sequences (Fig. 6I), further supporting the notion that the motifs identified in enhancers near down-regulated genes during JUND knock down may represent indirect changes in SP5 activity.

Experimental validation would be required to state with confidence whether differences in gene expression upon JUND knock down are more directly attributable to decreased SP5 expression than to changes in JunD expression on JUND knock down, as suggested by our motif analysis. Nevertheless, we find it intriguing that SP5 has been described as an activator and a repressor^30,31^, with^30^ explaining the discrepant observations regarding SP5 as an activator or repressor in part as a result of expression levels of SP5, with SP5 overexpression potentially causing it to behave as an activator in work by^31^, and endogenous levels of SP5 leading to its behavior as a repressor in work by^30^. We suggest that this explanation would be consistent with the pattern of enrichments we observed for the 5-mer motifs identified in JUND RNA-seq associated enhancers, where, if the motifs truly reflect presence of SP5 on enhancers, increased expression would lead to more SP5 occupancy at enhancers with more binding sites. Thus, based on our analysis of the JunD ChIP-seq and RNA-seq data, and our follow-up analysis of SP5 ChIP-seq data, we suggest that ShapeME can be a powerful tool for discovering motifs in DNA that undergird important biological signals of interest, even when those signals arise due to indirect effects.

## Discussion

We have developed ShapeME as a powerful and flexible motif inference workflow that enables the *de novo* discovery of short motifs in either DNA sequence space, DNA shape space, or both. A key advantage of our workflow is its sheer flexibility as it can discover motifs that are explanatory for diverse data types from simple binary binding data to complex multi-category expression patterns, and it tests the suitability of discovered sequence and shape motifs to only retain the minimum set of motifs needed for prediction. The flexibility of our algorithm enables us to capture the binding preferences of proteins that are primarily mediated through interactions with the Watson-Crick-Franklin face of the DNA (sequence) as well as binding preferences that are more generally driven by the properties of the DNA surface (shape) through a small number of biophysically interpretable parameters. Furthermore, our *de novo* shape motif discovery algorithm outperforms recently published state-of-the-art *de novo* shape motif algorithm ShapeMotifEM, with higher AUPR for discriminating between bound and non-bound sequences in ChIP-seq benchmarks.

We show the utility of using DNA shape information to predict binding for diverse DNA binding proteins in eukaryotes and bacteria. For most human proteins, DNA shape motifs alone or in combination with sequence motifs improve binding prediction. For bacterial chromatin proteins, explanatory sequence motifs have been elusive, reflecting the broad and apparently degenerate binding properties of these enigmatic proteins. Here, we show that the simple physical property of DNA roll may help explain the binding preferences of the abundant *E. coli* chromatin protein H-NS. Given H-NS’s role in silencing horizontally acquired genes, a broad recognition signature predicated on the DNA surface properties is likely beneficial to enable non-sequence specific binding of foreign DNA.

Since mutation of DNA binding sites primarily results in changes in base-pair content, protein recognition through non-Watson-Crick-Franklin surfaces of the DNA may enable some level of mutational tolerance to changes in DNA recognition sites; it is also notable that the structural properties of DNA are often robust to sequence changes, especially for GC rich regions (Fig. 1D-E). It is also possible that DNA surface properties are more sensitive to cellular state through changes in local DNA supercoiling or overall chromatin structure that is not captured in current predictions of DNA shape from sequence. At present, our work is limited by the reliance on a single idealized model for deriving shape parameters from primary sequence, which ignores cellular state, potential impacts on structure of nearby binding sites of DNA-binding proteins, and modifications such as DNA methylation. Future work incorporating the effects of cellular state on DNA shape may improve the detection and predictive utility of short shape motifs for DNA binding proteins. There is also a need for ongoing work to better develop a framework for comparing shape motifs with each other, with random background distributions, and with the known binding preferences of characterized DNA binding proteins. Unlike sequence motifs, it is much more difficult to engineer a given shape motif into a given sequence context. Tools to make targeted changes to sequence to shift binding preference for a given DNA binding protein whose binding preference is primarily driven by DNA shape features are currently lacking. Here we show that DNA shape motifs are broadly useful to predict protein binding but future work to make targeted changes in DNA shape space coupled with complementary amino acid changes in DNA binding proteins will give mechanistic insight into how DNA binding proteins use the entire DNA surface to select target sites.

## Materials and Methods

### ShapeME input data and arguments of note

The simplest user experience for ShapeME is for the user to prepare two input files, described below, and to submit them to the ShapeME web server using default parameters. All subsequent steps of shape motif inference and optional sequence motif inference are then automated by ShapeME. The two input files are:

- Score file

○ A tab-delimited file with a single header row
○ Header has columns “name” and “score”
○ Scores are usually binary or categorical
○ Scores can be continuous, in which case they will be quantized into *q* discrete categories prior to motif inference, where *q* is a value supplied by the user
- Sequence file

○ Fasta format. One DNA sequence for each record named in the “name” column of the score file.
○ All DNA sequence lengths must be identical and must be at least *k*+4 long, where *k* is the length of shape motifs to infer.

Arguments of note affecting the behavior of the ShapeME workflow include the following:

● Shape motif length (integer) - described briefly above as *k*, sets the shape motif length the shape-based component of ShapeME will return
● Find sequence motifs (boolean) - if True, ShapeME will still use its shape-based component to infer shape motifs, and will also use STREME to identify sequence motifs. Shape and sequence motifs will be separately pruned using CAMI and LASSO regression. If informative motifs of only a single type, i.e., if only shape motifs or only sequence motifs made it through their respective CAMI/LASSO filters, ShapeME returns the motifs of the remaining type to the user. However, if both sequence motifs and shape motifs made it through these pruning steps, they are merged into a single motif model and a final round of LASSO regression is performed.
● Number of quantiles to discretize continuous scores into (integer) - Described as *q* above, this is the number of quantiles into which ShapeME will bin sequences for motif inference.
● Sequence motifs positive categories (integer or list of integers) - If “Find sequence motifs” is True, this argument identifies the “positive set” of sequences for STREME. By default we assume the user has input binary scores and that sequences labeled with score 1 are the positive set.

○ If the user supplies categorical or continuous scores, they must choose which scores denote the positive set. For example, if the scores are integers from 1-5, inclusive, setting this argument to 5 will use sequences only in category 5 as the positive set. Setting it to “4,5” will include sequences in both categories 4 and 5 as the positive set.
○ For continuous input scores it is important to keep in mind that when ShapeME bins input sequences into *q* quantiles, the resulting categories are zero-indexed, with category 0 containing the lowest-scoring sequences and category *q*-1 containing the highest-scoring sequences. Therefore, if the highest-scoring sequences should be searched for sequence motifs by STREME, the user should set this argument to *q*.
● Maximum number of shape motif occurrences on each strand (integer) - By default the shape-based component of ShapeME counts up to 1 shape motif match per strand of a given query piece of DNA. The user may increase this number up to 4. Note: increasing this argument can dramatically increase ShapeME run times.

### General Description of the Algorithm

Automated shape motif inference is performed by the shape-based component of ShapeME in the following steps, explained simplistically here and discussed in greater detail below:

1. Produce input shape array *A*.

a. Convert input sequences to shape parameters EP, HelT, MGW, ProT, and Roll using DNAShapeR^9^.
b. Standardize resulting shapes to z-scores by considering the average and standard deviation of each shape parameter for all sequences within the dataset.
2. Identify a reasonable starting threshold for Manhattan distance, under which a comparison between a query and reference will be considered a “hit”.
3. Screen batches of *S* in *A* for informative seeds.

a. By default, fetch 500 *S* for each batch
b. Given the user-selected window size, *k*, compare each *k*-width window, *b*, in every *S* in this batch, to every other *b* in *A* to identify hits between *b* and each *S* in *A*.
c. Calculate adjusted mutual information (AMI) between *b* hits and Y.
d. Sort all *b* in this batch in order of descending AMI. Retain the *b* with highest AMI, and any additional *b* that adds information to the model in addition to that provided by prior seeds, including prior seed in this batch and any prior batch, as judged by conditional adjusted mutual information (CAMI).
e. If informative seeds were identified in step 3c and more batches of sequences are available to search for more seeds, move to step 3A. If informative seeds were identified in step 3c and no more sequences are available to search for seeds, move to step 4. If no informative seeds were identified in 3c, retain the top 5 low-information seeds and move to the next batch of sequences if available. If no more input sequences are available, move to step 4.
f. By default, if five batches have been searched for seeds without any new informative seeds entering the pool, end the search and move to step 4.
4. Optimize shape, weight, and “hit” threshold values via maximizing AMI between each motif’s hits and input scores.
5. Sort motifs in order of descending AMI and prune motifs, retaining only those that add information in addition to prior motifs as determined by CAMI.
6. Perform model selection using LASSO regression, using the input scores as the outcome variable. The design matrix is composed of a column representing the intercept and an additional column representing each motif’s hit status for each row’s input *S*.
7. Remaining motifs are written to a “dsm” file, which is MEME-like in format. If only the intercept remains after LASSO regression, no informative shape motifs were identified and no “dsm” file is produced.
8. Each motif’s enrichment in the input categories is calculated and a heatmap is prepared to summarize motif enrichment.
9. Motif logos are plotted.
10. Steps 1-10 are performed for each of 5 folds of the input data and cross-validated performance measures are calculated and plotted along with performance of the model trained on all data in the input dataset.

### Application of information theory to shape-based motif inference

Adjusted mutual information (AMI) is defined as in equation 27 of Vinh and colleagues^32^:

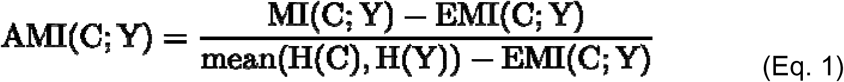

Where MI refers to mutual information,

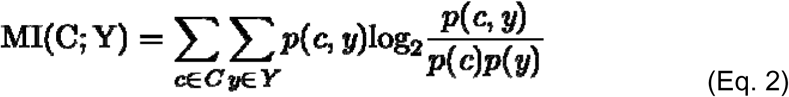

H refers to information entropy,

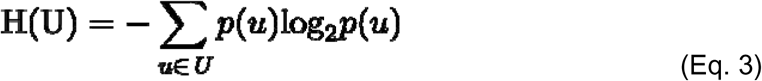

where in equation 3 above we use the term U to represent any input vector of category assignments with support set *U*. EMI in equation 1 denotes expected mutual information, which is the mutual information expected by random chance when permuting over all possible category assignments for a pair of vectors as defined in equation 24a by Vinh and colleagues^32^.

Pruning of redundant motifs from a set of motifs is performed through a combination of conditional adjusted mutual information (CAMI) and LASSO regression. CAMI is defined as follows:

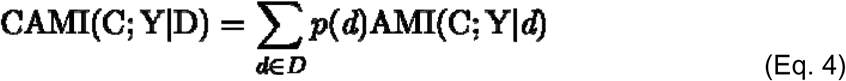

where D represents a hits vector for a prior, more informative motif than the motif represented by hits vector C. Hits vector D has support set *D*.

### Pruning of motifs based on CAMI

We use an AIC-like information criterion, which we refer to as the Information Content of motif *m* (IC*_m_*), to define whether a motif’s information content is sufficient to include it in the set of motifs reported by ShapeME. The criterion balances information content of the motif under consideration for addition to the model with the number of parameters added to the overall motif model:

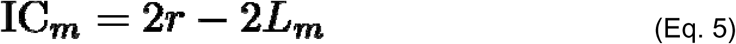

where IC*_m_* is the information criterion for motif *m* and *L_m_* is the log-likelihood for motif *m*. Our definition of *L_m_*is:

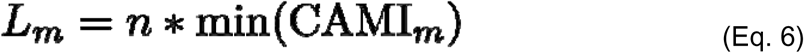

where *n* is the number of sequences in the input dataset. Here we note *L_m_* arises from the minimal CAMI for motif *m* compared against all motifs with higher AMI already present in the model. In practice, this ensures that if any motif already present in the model provides similar information as *m* so as to make adding motif *m* to the model (along with its extra *d* parameters) insufficiently informative, then *m* is not included in the final model.

### Determination of motif matches

We use a weighted Manhattan distance as the measure of “closeness” of DNA shapes. The measure is defined below:

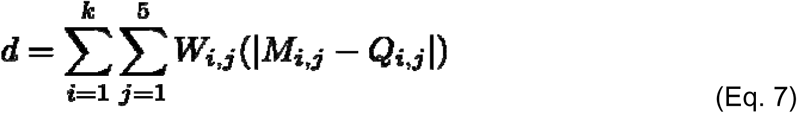

In order to determine a weighted Manhattan distance value under which a seed is considered a “hit” against a query shape, an initial threshold for the distance measure is found by sampling 500 random shape *k-*mers from the input sequences. Each randomly selected seed is compared to the forward and reverse strands of two randomly-chosen *k-*mers within each input sequence. The initial threshold is calculated to be the mean of all resulting distances minus two standard deviations.

For each seed or motif, for every *S* in *A*, each *Q k*-mer is compared to the seed or motif and the number of hits to distinct *Q k*-mers on each strand of *S* is counted up the the maximum allowed hit count per strand.

Hits arrived at by the above approach are used to construct a hits vector, C, for each seed or motif. Each strand is searched for hits to each seed or motif. If a motif matches once on each strand its hit value is assigned “1,1”. If it matches once to a single strand, regardless of which, its hit value is “0,1”. A comparison with no matches results in a hit value of “0,0”.

### Shape motif optimization

After identifying an informative set of seeds, the values in each seed’s *M* and *W* matrices, and each seed’s *t* are optimized using a simulated annealing approach, after which the seeds are now termed “motifs”. By default, 20,000 iterations are performed. For each iteration of optimization, a random parameter, *p*, where “parameter” here refers to any value of *M*, *R*, or *t* is nudged from its prior value by a value randomly sampled from a normal distribution of mean 0.0 and standard deviation 0.25. The updated value of *p* in iteration *i* of the optimization procedure is:

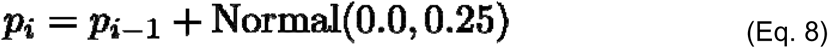

If *p* was selected from either *M* or *R* we do not allow *p_i_* to take values less than -4.0 or greater than 4.0, and if the above perturbation violates those constraints, we manually re-set *p_i_* to either -4.0 or 4.0, respectively. Similarly, perturbed values of *t* are constrained to be between 0.0 and 3.0. If *p* was selected from *R*, we update *W* as follows:

First we apply a constrained inverse-logit transformation to each value in *R* such that transformed values (*R*^✝^) range from a lower limit of α and an upper limit of 1.0:

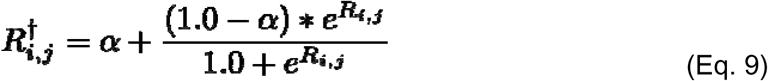

We then calculate *W* to be:

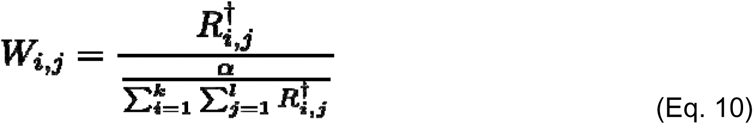

After perturbing *p* and updating *W* if appropriate, the updated motif’s AMI is calculated. The objective of optimization is to maximize AMI, and updates resulting in increased AMI are always accepted, but those resulting in decreased AMI are accepted with the following probability, *p*(*a*):

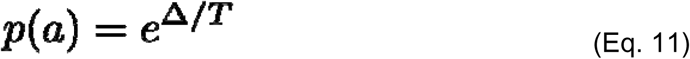

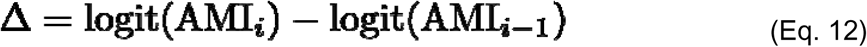

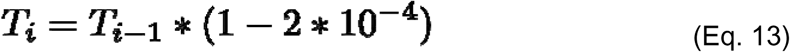

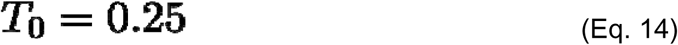

where *i* in equations 12 and 13 refers to a given iteration of optimization, and *T* in equations 11, 13, and 14 represent a “temperature” parameter which is initialized to be 0.25 (equation 14) and is decreased by a small fraction each iteration (equation 13). By default, ShapeME performs 20,000 iterations of simulated annealing. After each motif has been optimized in the above manner, formerly distinctively-informative motifs may have converged to now provide similar information content, so we prune the optimized motifs as described in section “Pruning of motifs based on CAMI”.

### Incorporation of sequence motifs using STREME and FIMO

If a user selects to use the sequence-based component of ShapeME, sequence motifs enriched in the user-defined “positive set” of sequences (see section “ShapeME input data and arguments of note” above for details on defining the positive set of sequences) are identified using STREME version 5.5.3. Any sequence motifs returned by STREME with an e-value less than 0.05 are retained for further evaluation. For sequence motifs, “hits” are identified using FIMO, where a sequence matching a motif with a q-value less than 0.05 receives a 1 in the motif’s hits vector and all other sequences receive a 0. The binary hits vector for each sequence motif is used for the purposes of AMI and CAMI calculation and LASSO regression. Note that CAMI-based pruning of sequence motifs and a first round of LASSO regression are performed separately from CAMI-based pruning and LASSO regression for shape motifs. If both sequence and shape motifs are to be identified by ShapeME, a final round of LASSO regression is performed for final model selection after merging filtered shape and sequence motifs into a single model.

### Model selection with LASSO regression

To select a final set of informative motifs and hits categories, we used LASSO regression as follows. We set up a design matrix with an intercept column and a column for every motif and non-zero hit category. For example, in a model with two shape motifs and a maximum number of hits per strand of 1, the design matrix would have 5 columns (intercept, motif X hit category “0,1”, motif X hit category ”1,1”, motif Y hit category “0,1”, and motif Y hit category “1,1”). Each row of the design matrix represents a sequence in *A*. For each sequence motif, a single column is added to the design matrix to denote which rows (sequences) were hits to the given sequence motif. A LASSO regression model is fit to regress the input categories, Y, against the design matrix using the “cv.glmnet” function from the R package glmnet v 4.1-7. Arguments passed to cv.glmnet were alpha = 1, folds = 10, and family = “binomial” if Y is a binary vector or family = “multinomial” if Y is categorical. If any resulting coefficient was zero after fitting the model, its corresponding motif and hit category pair was removed from the overall motif model. This process often removes single hit categories for any given motif in the model and sometimes removes all hit categories for a given motif.

### Assessment of motif model performance

We used the area under the precision-recall curve (AUPR) to assess ShapeME motif model performance. Hits to sequence and shape motifs were identified, and predictions of which category each test sequence belonged to were made using the fitted coefficients from the LASSO regression step during motif inference (see section “Model selection with LASSO regression”). Precision and recall are calculated at several thresholds, and AUPR was calculated using the R package PRROC, version 1.3.1^33–35^.

### Analysis of 5-mer shape robustness

#### Sequence preparation and quantile rank difference

Every sequence 5-mer, flanked on each end by an “AA” dinucleotide, was prepared and converted to shapes using DNAShapeR^9^. The edit distance and Manhattan distance was calculated for every pair-wise comparison of 5-mer sequence and shape, respectively. We calculated the quantile of each shape or sequence distance against all other shapes or sequences by using the rankdata function in the scipy (version 1.11.1) module and dividing the result by the total number of distances. The difference in quantile rank between shape and sequence was simply each shape quantile minus the corresponding sequence quantile.

#### Shape robustness

Here we define the term “close-mer” to be any two sequences with an edit distance equal to 1. A given piece of DNA’s shape robustness is defined as its mean median-adjusted shape distance between all “close-mer” sequences. For median adjustment we took the global median close-mer shape distance, *g*, to be the median shape distance between all close-mers. To calculate shape robustness for a given 5-mer (flanked by “AA” on each side), *i*, we select all close-mers, *K*, to *i*. Robustness for sequence *i*, γ*_i_*, is then calculated as the mean median-adjusted shape distance between *i* and all of the close-mers of *i*, *K*:

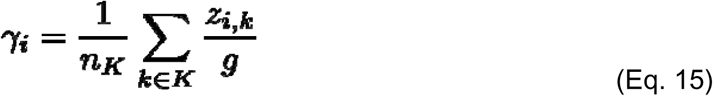

where *n_K_* is the number of close-mers to *i*, and *z_i,k_* is the shape Manhattan distance between *i* and its given close-mer *k*.

### *E. coli* sequencing data analysis

Accessions for data used in the *E. coli* motif analysis are given in Table 2.

**Table 1.** Symbols used in ShapeME description.

| Symbol | Description | Type | Shape |
| --- | --- | --- | --- |
| $n$ | Number of input sequences | Integer | NA |
| $l$ | Length of each input sequence after shape conversion | Integer | NA |
| $Y$ | Vector of categorical scores associated with each input sequence with support set $Y$ | Vector | $n$ |
| $A$ | The array of all input shape matrices | Array | $n \times l \times 5$ |
| $S$ | A shape matrix arising from a single input sequence of length $L$ | Matrix | $l \times 5$ |
| $k$ | Length of motifs to be identified | Integer | NA |
| $M$ | A $k$-mer shape matrix | Matrix | $k \times 5$ |
| $W$ | A $k$-mer weights matrix | Matrix | $k \times 5$ |
| $R$ | Matrix used to derive $W$ for each motif, used during motif optimization. | Matrix | $k \times 5$ |
| $e$ | A seed, defined by a shape matrix $M$, a uniform weights matrix $W$, and a hit threshold. | NA | NA |
| $m$ | A motif, defined by its shape matrix $M$, weights matrix $W$, and hit threshold. | NA | NA |
| $q$ | The user-selected number of quantiles that ShapeME will quantize continuous input data into. Binary and categorical input data are handled as-is. | Integer | NA |
| $d$ | The weighted Manhattan distance between a seed or motif and a query shape $k$-mer | Float | NA |
| $t$ | Threshold value below which a given $d$ is considered a “hit”. Each motif in a shape motif model has its own $t$ after optimization. | Float | NA |
| $C$ | The vector of hit classes for a given motif, with support set $C$. For each input sequence, a motif can match zero times, once, or potentially multiple times on each strand. The default behavior is to match up to one time on each strand. In this case, the possible hits classes are zero hits for a sequence (0,0), a hit on a single strand (0,1), or a hit on each strand (1,1). | Vector | $n$ |
| $r$ | Number of parameters in the overall motif model. Each shape motif adds $2 \times k \times 5 + 1$ parameters ($2 \times k \times 5$ arises because both an $M$ and a $W$ matrix define a motif. 1 is added because the motif's threshold for calling “hits” is also a parameter of each motif). Each sequence motif adds $3 \times \text{sequence-motif-length}$ to the overall model. | Integer | NA |

**Table 2.** *E. coli* H-NS data used in this work.

| SRA Accession | Description | Reference |
| --- | --- | --- |
| ERR015955 | H-NS ChIP-seq from *E. coli* in transition to stationary (TS) phase | Kahramanoglou and colleagues, 2011, DOI: 10.1093/nar/gkq934<sup>36</sup> |
| ERR015956 | H-NS ChIP-seq from *E. coli* in stationary (S) phase |  |
| ERR015957 | H-NS ChIP-seq from *E. coli* in early-exponential (EE) phase |  |
| ERR015958 | H-NS ChIP-seq from *E. coli* in mid-exponential (ME) phase |  |

#### H-NS data analysis

Single-end sequencing reads were aligned to the *E. coli* K12 MG1655 reference genome (GenBank ID: U00096.3) using bowtie2, version 2.4.4, with the default arguments^37^. The resulting SAM files were converted to BAM files and sorted using samtools, version 1.9, with default arguments^38^. Coverage was calculated using the following code:

> bedtools bamtobed -i {input_bamfile} \

> | bedtools genomecov -bga -i - -g {genome_file} > {ouput_file}

with bedtools, version 2.31.1^39^. Coverage values were converted to robust z-scores using the command “bgtools robust_z” from our bgtools package (https://github.com/jwschroeder3/bgtools).

The *E. coli* genome was divided into either 100 or 300 bp windows, and the median robust z-score was mapped to each window using bedtools. The sequence of each window and its paired median robust z-score of H-NS occupancy were submitted to ShapeME in shape-only mode to identify shape motifs informing H-NS occupancy. ShapeME was set to quantize the sequences into deciles based on their H-NS occupancy z-scores.

### ShapeME analysis of ENCODE data

#### Data retrieval and preparation for ShapeME

We downloaded bed files containing irreproducible discovery rate thresholded peaks for 29 transcription factors from ENCODE (see Table 3 for factors and ENCODE file IDs). To prepare sequence and score files from ENCODE datasets for motif inference we ran the prep_data subcommand of ShapeME as follows:

> singularity exec -B {data_directory} shapeme.sif \ python /src/python3/ShapeME.py prep_data \

> --fasta_file GCF_000001405.26_GRCh38_chr_chroms.fa \

> --data_dir {data_directory} \

> --narrowpeak_file {encode_file_id}.bed \

> --wsize 60 --nrand 3 --max_peaks 1000

where `--wsize 60` directs prep_data to write 60 base pair sequences from the geometric center of each peak or randomly selected genomic location, `--nrand 3` causes three times as many random genomic loci to be selected than peaks, and `--max_peaks 1000` causes datasets with greater than 1000 peaks to down-sampled such that 1000 peaks are randomly selected for motif inference. The prep_data subcommand automatically sampled peaks from the input bed file, extracted peak and random genomic loci sequences from the input fasta file, and wrote sequences and scores to their appropriate files.

**Table 3.** ENCODE and JASPAR identifiers for datasets used in this work.

| TF name | ENCODE experiment ID | ENCODE file ID | JASPAR motif ID |
| --- | --- | --- | --- |
| ETS1 | ENCSR681WHQ | ENCFF191NVV | MA0098.3 |
| YY1 | ENCSR859RAO | ENCFF430PDB | MA0095.2 |
| SRF | ENCSR000BLV | ENCFF050QMU | MA0083.3 |
| HNF4G | ENCSR000BNJ | ENCFF123FQW | MA0484.1 |
| ELF1 | ENCSR000BMZ | ENCFF839JEO | MA0473.2 |
| ZEB1 | ENCSR000BVN | ENCFF060IPX | MA0103.3 |
| ZBTB33 | ENCSR000BHR | ENCFF787GZD | MA0527.1 |
| TEAD4 | ENCSR800JRG | ENCFF355HRT | MA0809.1 |
| TCF3 | ENCSR000BQT | ENCFF700TAS | MA0522.2 |
| SP4 | ENCSR000BQV | ENCFF257FUV | MA0685.1 |
| SP2 | ENCSR000BOU | ENCFF591XXC | MA0516.1 |
| SP1 | ENCSR334KIQ | ENCFF594LBX | MA0079.3 |
| RUNX3 | ENCSR000BRI | ENCFF346JDW | MA0684.1 |
| POU5F1 | ENCSR364SNE | ENCFF724DNU | MA1115.1 |
| POU2F2 | ENCSR000BGP | ENCFF934JFA | MA0507.1 |
| PBX3 | ENCSR000BTN | ENCFF656HFI | MA1114.1 |
| NR2F2 | ENCSR000BVM | ENCFF804MKP | MA1111.1 |
| NFIC | ENCSR000BQX | ENCFF611FIV | MA0161.2 |
| MEF2C | ENCSR000BNG | ENCFF238UKB | MA0497.1 |
| MEF2A | ENCSR000BNV | ENCFF031NTF | MA0052.3 |
| MAX | ENCSR000BTM | ENCFF104FEX | MA0058.3 |
| JUND | ENCSR000EEI | ENCFF562YSV | MA0491.1 |
| IRF4 | ENCSR000BGY | ENCFF113VGD | MA1419.1 |
| GATA2 | ENCSR000BKM | ENCFF242YZU | MA0036.3 |
| FOSL1 | ENCSR000BTE | ENCFF936WPZ | MA0477.1 |
| EBF1 | ENCSR000BGU | ENCFF895MHN | MA0154.3 |
| E2F6 | ENCSR000BTC | ENCFF786NAO | MA0471.1 |
| FOXA1 | ENCSR267DFA | ENCFF011QFM | MA0148.3 |
| CTCF | ENCSR000DWH | ENCFF948CYD | MA0139.1 |
| SP5 | ENCSR019NPF | ENCFF347UBA | NA |

To prepare datasets for benchmarking ShapeME against ShapeMotifEM, we included the peaks above the 95th percentile of signal strength by invoking the prep_data subcommand as follows:

> singularity exec -B {data_directory} shapeme.sif \

> python /src/python3/ShapeME.py prep_data \

> --fasta_file GCF_000001405.26_GRCh38_chr_chroms.fa \

> --data_dir {data_directory} \

> --narrowpeak_file {encode_file_id}.bed \

> --wsize 60 --nrand 3 --percentile_thresh 0.95

This call includes all peaks above the 95th percentile in the output fasta and scores files.

#### ShapeME runs

ShapeME was run in “both” mode for every dataset using the following basic command line invocation:

> singularity exec -B {data_directory} shapeme.sif \

> python /src/python3/ShapeME.py infer \

> --find_seq_motifs --data_dir {data_directory} \

> --seq_fasta seqs.fa --score_file seqs.txt \

> --crossval_folds 5 --nprocs 64 --alpha 0.01 \

> --max_count 1 --temperature 0.25 --t_adj 0.0002 \

> --opt_niter 20000 --stepsize 0.25 --threshold_constraints 0 3 \

> --shape_constraints -4 4 --weights_constraints -4 4 \

> --batch_size 500 --max_batch_no_new_seed 5 \

> --kmer 10

For inference of only shape motifs, the call was modified to omit the `--find_seq_motifs` flag. For inference of only sequence motifs, the call included both the `--find_seq_motifs` flag and the `--no_shape_motifs` flag.

To perform sequence and shape inference where the sequence motif was from the JASPAR database, the call was similar to above with the sole modification being the addition of `--seq_meme_file jaspar_motifs.meme` to the call. See Table 3 for the JASPAR motif ID for each transcription factor.

#### Statistical analysis of 5-fold cross validated ShapeME results for assessing differences in performance between shape, sequence, and both modes

To state which set of ShapeME results performed best for each ENCODE dataset in terms of Bayesian evidence ratios (K), we performed the Bayesian analog of the t-test, Bayesian estimation supersedes the t-test (BEST^40^). For each transcription factor, we fit the BEST model to the 5 AUPRs from 5-fold cross validation for each of two ShapeME model types. K was calculated to be the number of posterior samples of the difference in the means between the two model types above zero divided by the number of posterior samples below zero. We applied a threshold of K > 3 to consider a motif model to perform better than the competing model. That is to say, greater than three times as many samples of the posterior distribution of the difference in the inferred mean cross-validated performance had to support one model outperforming the other for the threshold to be passed. The BEST model was fit using pymc^41^; code and input AUPRs can be found at our github repository (https://github.com/jwschroeder3/shapeme_best_regression).

#### Calculating area under precision-recall curves to assess ShapeMotifEM performance

We compiled ShapeMotifEM within an Apptainer container and trained shape motif models using the five shape parameters EP, HelT, MGW, ProT, and Roll using the R script “learn_motifs.R” found at the github repository https://github.com/jwschroeder3/shapemotifem_analysis_code.git. The basic command line call for each transcription factor was:

> singularity exec -B {data_directory} shapemotifem.sif \

> Rscript /src/shapemotifem_analysis_code/learn_motifs.R \

> -f {training_sequences.fa} \

> -d {output_directory}

The training sequences for every ShapeMotifEM run were identical to those used to train ShapeME motif models, except that ShapeMotifEM is designed to learn motifs enriched within a single “positive” set of sequences. Therefore, we selected only the “peak” sequences from each ShapeME training dataset for use in training ShapeMotifEM models. The test sequences used to evaluate ShapeMotifEM runs on entire datasets or each of 5 folds performed for 5-fold cross validation were identical to those used to evaluate ShapeME models.

To evaluate ShapeMotifEM performance, we first converted test sequences with known “positive” or “negative”, i.e., peak or non-peak, respectively, to z-scores and used the mean and standard deviation for the test sequences to convert the shape values for the motifs learned by ShapeMotifEM to z-scores on the same scale. We compared every standardized test sequence to each standardized motif returned by ShapeMotifEM by sliding the motif along each test sequence and calculating the Manhattan distance for each comparison. We retained the minimum Manhattan distance across all motifs and windows for each test sequence. We next called test sequences as “positive” or “negative” at several Manhattan distance thresholds and calculated the precision and recall for each threshold value. The area under the precision-recall curve was calculated using the trapezoidal rule. Our code for performing the shape value standardization, Manhattan distance calculation, and AUPR calculations can be found in the github repository (https://github.com/jwschroeder3/shapemotifem_analysis_code.git).

### JunD knock down differential expression and ShapeME analysis

#### Differential expression

CRISPRi-mediated knock down RNA-seq result tsv files were downloaded for the non-specific guide control samples ENCFF742HVE and ENCFF465RMN, and for the JUND-targeting guide samples ENCFF068OFG and ENCFF725VDN^22–25^. The “expected_count” column was retained for each gene and used to prepare a counts matrix for differential expression analysis using DESeq2 version 1.30.1^42^. Genes included in the analysis were those for which at least two samples had at least 5 counts each. The DESeqDataSet was set up to fit an intercept and a term for the “KD” condition. The contrast tested was KD/control, and shrunken log_2_(fold-change) in transcript abundance was calculated using the lfcShrink function from DESeq2, with argument “type” set to “ashr”^43^. Code and data used for our differential expression analysis can be found at the following github repository (https://github.com/jwschroeder3/src_for_JunD_KD_analysis.git).

### ShapeME analysis of JunD knock down results

#### Associating enhancers with gene shrunken log_2_(fold-change)

We downloaded the annotated regulatory regions and genes from BioMart^44^ for human reference sequence GRCh37.p13. We extracted only enhancers from the regulatory region annotations and converted the gene annotations to bed format and enhancer annotations to gff format. To identify the closest enhancer to each gene we used bedtools version 2.31.1:

> bedtools closest -a {biomart_genes.bed} -b {enhancers.v112.gff} -d -t

> first > closest_enhancer_to_each_gene.bed

Note that using the above command, if multiple enhancers are found within a gene, i.e., their distance from the gene is zero and are thus tied as the closest enhancer to the gene, the first enhancer in genome coordinates is selected as the “closest” enhancer to the gene. We removed genes for which the closest enhancer was greater than 450,000 base pairs from the gene. To associate enhancers with the shrunken log_2_(fold-change) values inferred above, we merged the DESeq results tables to the remaining genes by joining on gene identifier. Because ShapeME requires all input sequence lengths to be identical, we next set each enhancer’s coordinates to be the central 100 base pairs of the original annotation. We converted these results to a bed file and extracted the enhancer sequences using bedtools getfasta:

> # Edit fasta headers to match chromosome identifiers in enhancer

> # annotations

> sed ’s/^>NC_0\+/>/g’ GCF_000001405.25_GRCh37.p13_genomic.fna \

> > edited_GRCh37.fa

> sed -i ’s/\.[[:digit:]]\+ Homo/ Homo/g’ edited_GRCh37.fa

> sed -i ’s/^>23/>X/g’ edited_GRCh37.fa

> # get sequences of enhancers bedtools getfasta \

> -bed middle_100pb_enhancers.bed \

> -fi edited_GRCh37.fa \

> -fo enhancer_sequences.fa \

> -nameOnly

For the binarized input scores used in Fig. 6B-C we set all genes with shrunken log_2_(fold-change) >= 0 as category 1, and all other genes as category 0. For the categorical input scores used in Fig. 6D-E we passed the continuous shrunken log_2_(fold-change) scores to ShapeME with *q* set to 5.

#### ShapeME commands

ShapeME run on binary input scores in “both” mode to infer 10-mer shape motifs and any possible sequence motifs was performed with the following command line invocation:

> singularity exec -B {data_directory} shapeme.sif \

> python /src/python3/ShapeME.py infer \

> --data_dir {data_directory} \

> --seq_fasta {seq_file.fa} --score_file {score_file.txt} \

> --find_seq_motifs --streme_thresh 0.05 --seq_motif_positive_cats 1

> --crossval_folds 5 --kmer 10 --max_count 1 \

> --threshold_sd 2 --init_threshold_seed_num 500 \

> --init_threshold_recs_per_seed 100 \

> --init_threshold_windows_per_record 2 \

> --nprocs 64 --threshold_constraints 0 3 \

> --shape_constraints -4 4 \

> --weights_constraints -4 4 \

> --temperature 1 --t_adj 0.01 --stepsize 0.25 \

> --opt_niter 20000 --alpha 0.1 \

> --max_batch_no_new_seed 5 --batch_size 500 \

> --max_n 100000 --log_level info

Using continuous input scores to search only for 5-mer shape motifs with a maximum count per strand of 2 was performed as above, with the following modifications to the call. The input score file contained the continuous shrunken log_2_(fold-change) values instead of binarized values, `-- find_seq_motifs` was omitted, `--kmer` was set to 5, and `--max_count` was set to 2.

#### ShapeME analysis of SP5 ENCODE data

The following call was used to run ShapeME to search for shape motifs in SP5 ChIP-seq peaks:

> singularity exec -B {data_directory} shapeme.sif \

> python /src/python3/ShapeME.py infer \

> --data_dir {data_directory} \

> --seq_fasta {seq_file.fa} --score_file {score_file.txt} \

> --crossval_folds 5 --kmer 10 --max_count 1 \

> --alpha 0.01 --temperature 0.25 --t_adj 0.0002 \

> --opt_niter 20000 --stepsize 0.25 \

> --threshold_constraints 0 3 --shape_constraints -4 4 \

> --weights_constraints -4 4 --batch_size 500 \

> --max_n 15000 --max_batch_no_new_seed 5

In contrast to other ENCODE datasets used in this work, we allowed a maximum of 15,000 sequences to be used as inputs for this ShapeME run. ShapeME automatically handled random selection of input sequences to prune the original input data (which contained almost 100,000 sequences of SP5 peaks and random genomic loci) to 15,000 sequences.

## Acknowledgments

This work was supported in part by the National Institute of General Medical Sciences, National Institutes of Health grant R35 GM128637 to L.F. In addition this work was supported by the National Science Foundation Graduate Research Fellowship DGE1256260 awarded to M.B.W.

## Author contributions

Michael Wolfe (Initial algorithm design and author, data analysis, writing), Jeremy Schroeder (algorithm optimization and refinement, data analysis, writing), Lydia Freddolino (conceptualization, supervision, funding, data analysis, writing)

---

**Supplementary Figure 1.**
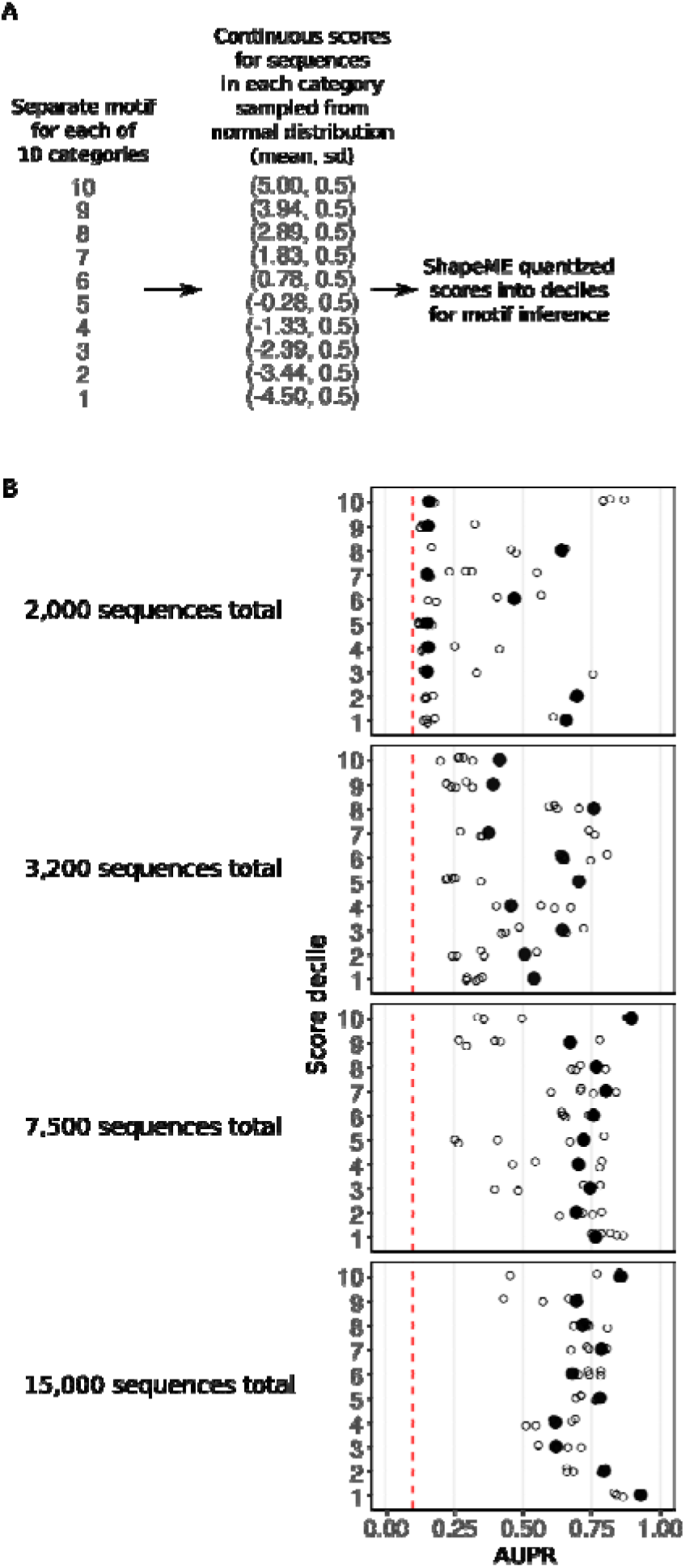
Effect of increasing record number on ability of ShapeME to detect shape motifs. A) Description of how continuous scores were generated for input sequences. Each category represents a set of sequences with one motif, present in each sequence once on each strand. ShapeME later quantized the input scores into deciles for motif inference. Note the relatively wide standard deviation for each category’s distribution, which makes this a very challenging task without sufficient data to inform motif inference. B) Area under precision-recall curves for each decile, at the indicated number of sequences in each synthetic dataset. The red dashed line denotes the performance that would be achieved by random chance.

**Supplementary Figure 2.**
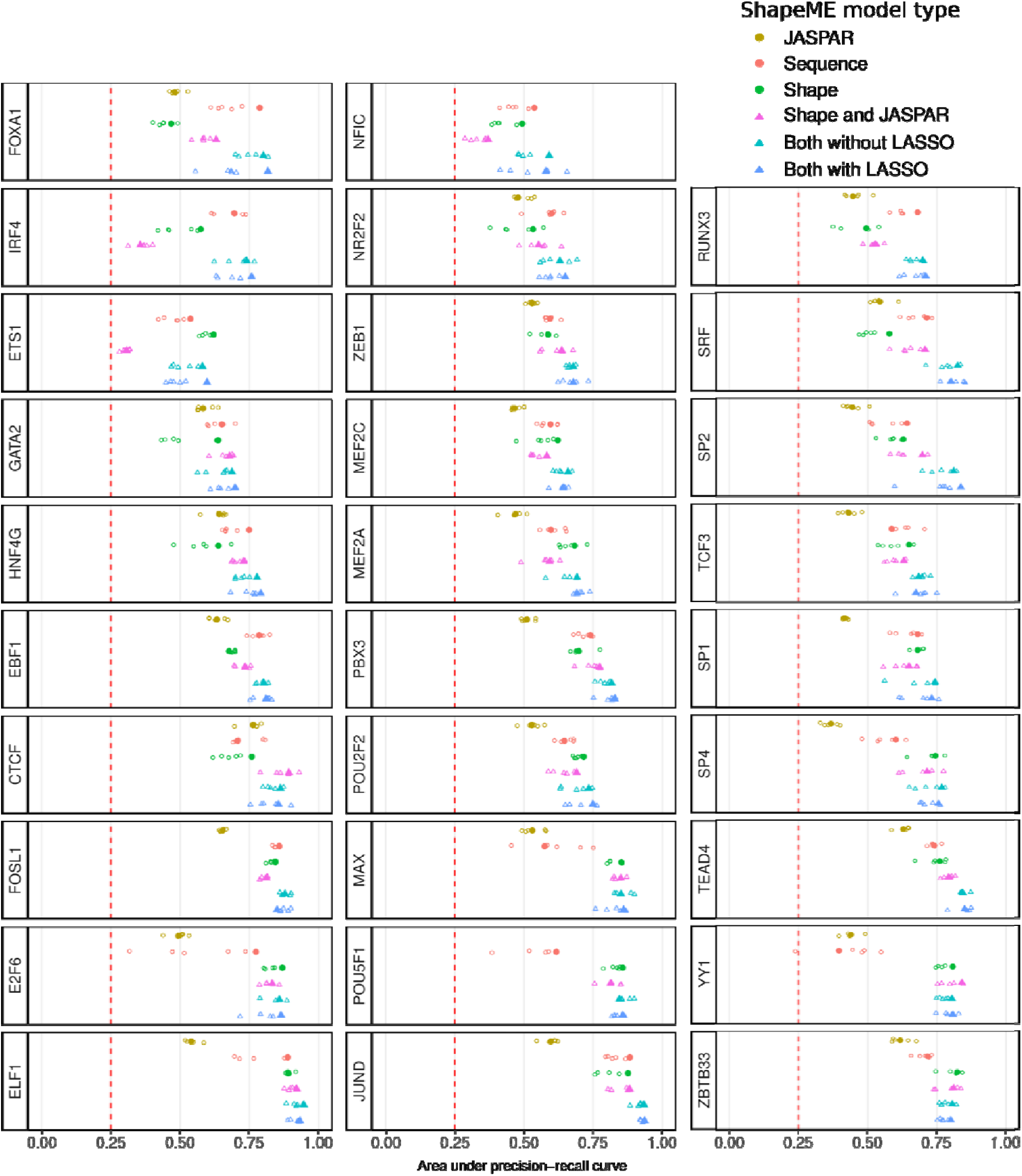
ShapeME performance in each of the modes denoted by marker colors, on each set of transcription factor peaks. 1,000 randomly selected peaks were taken from the irreproducible discovery rate passing peaks available through the ENCODE portal. Table 3 provides details on ENCODE datasets used.

## Notes

https://seq2fun.dcmb.med.umich.edu/shapeme/

### Competing Interest Statement

The authors have declared no competing interest.

